# Group-level phenotypes are idiosyncratic yet unpredictable over development in a clonal fish

**DOI:** 10.64898/2026.02.03.700030

**Authors:** Ammon Perkes, Jolle Jolles, Carolina Doran, David Bierbach, Max Wolf, Kate L Laskowski

## Abstract

Behavioral differences among groups can have profound impacts on individual fitness^1^, yet how group-level phenotypes arise and change during development is poorly understood. Group-level differences may originate from genetic differences or subtle variation among individuals^2–5^, but they can also emerge over time through social feedback mechanisms within and between groups, potentially producing dramatically different group-level behavior^6–8^. Disentangling these drivers of group phenotypes is challenging because groups are typically studied after they are already well established. Using the naturally clonal Amazon molly, we establish replicate groups of genetically identical individuals and continuously track group-level behavior from birth throughout the first 55 days of life. This design allows us to control for genetic and environmental sources of variance, thereby establishing a null baseline for group-level differences. Furthermore, by recording over an extended period beginning from day 1 of life, we test whether group phenotypes follow predictable trajectories driven by individual differences or instead diverge due to social feedback during development. We find that average behavioral changes across groups follow predictable patterns during this early developmental phase. In contrast, differences among groups are highly unstable over longer timescales: although groups differ from very early in life, it appears that the rank-order of group-level phenotypes undergoes a shuffling mid-development, around 4-5 weeks of age. These results suggest that distinct processes generate group-level variation across ontogeny, with early differences largely reflecting consistent individual variation; then as individuals mature and become more socially responsive, social feedbacks push groups along divergent, less predictable behavioral trajectories.

## Results & Discussion

Group living is common across organisms, ranging from transient aggregations to stable groups with complex sociality. In many taxa, including fishes, social attraction emerges within weeks of birth and continues to intensify until individuals actively maintain cohesive groups^9–14^. Many studies have explored the mechanisms and consequences of living in groups^15^ but despite apparently common rules governing social attraction, groups often differ in key group-level phenotypes, including tasks such as foraging, rearing offspring, or navigation^6,16–19^. These group-level differences appear to stem from a range of factors including group size^20–22^, composition^1,2,23^, and environmental context^24–26^. Yet these factors alone do not fully explain how group phenotypes arise, because social interactions can themselves generate feedback that shapes collective behavior^6,17,27^. While differences in group-level behavior have been studied extensively, often the groups studied have been established for multiple generations and can differ in number, age distribution, and environmental location, making it difficult to disentangle the effects of individual differences from social interactions. We lack experiments that follow the emergence of group-level phenotypes in replicate groups under controlled conditions from their formation^25,28,29^.

Although group-level phenotypes have not yet been tracked throughout development, existing research suggests that differences would likely be present from birth. Recent work has demonstrated phenotypic variation among clonal individuals raised under identical environments^30–35^. Such individuality can emerge very early in development and often phenotypes continue to diverge as individuals age^31,32,35–37^. Given this, groups may be expected to exhibit stable phenotypic differences reflecting the averaged behavior of their constituent members^38^ (Figure 1A, Figure S1). Alternatively, social interactions among group members may amplify or diminish initial differences, pushing groups along divergent^17,18^ (Figure 1B) or convergent^25^ (Figure 1C) developmental trajectories. Under each of these three hypotheses, group-level differences are largely predictable over time, but this need not be the case: if complex individual–group interactions^39^ operate alongside rapid developmental changes in physiology and behavior^12,40–42^ (Figure 1D), we might expect group phenotypes to lack long-term predictability, as the individual attributes and even the rules which determine group behavior could vary over time. Distinguishing among these alternative developmental hypotheses and identifying the mechanisms that could explain these dynamics requires datasets with high temporal resolution that follow directly comparable groups from their formation onward.

**Figure 1.**
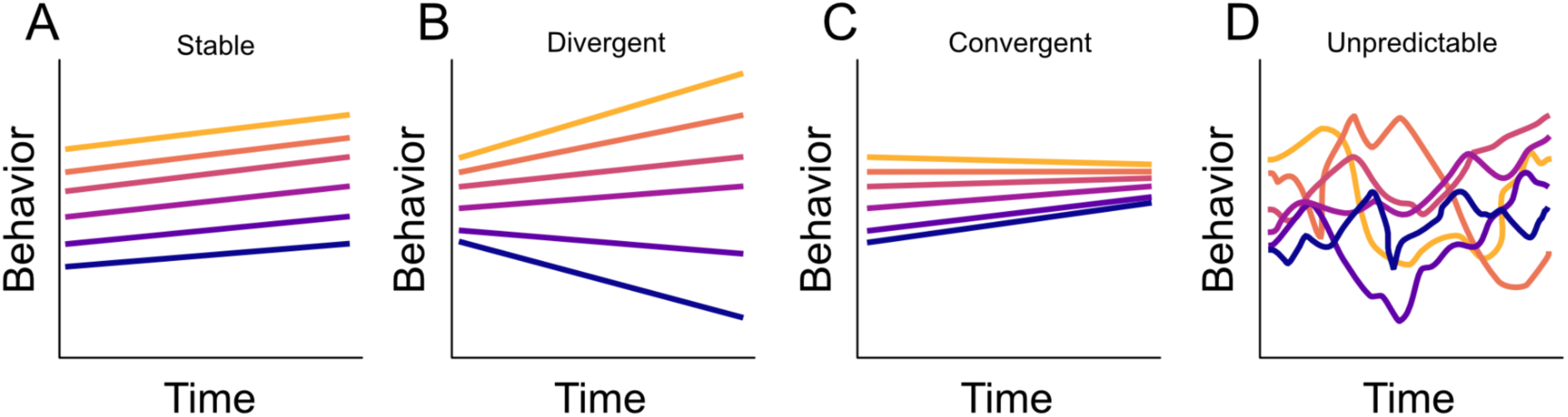
Four possible trajectories of group-level behavior over time. Each line represents the group phenotype (e.g., average velocity). In A) Among-group differences are consistent over time. In B) differences increase, whereas in C) among-group differences diminish over time. In D) group-phenotypes are unpredictable over development.

Here, we take advantage of the naturally clonal Amazon molly (*Poecilia formosa*) to track the emergence of group phenotypes continuously from birth. Amazon mollies are a live-bearing fish species without parental care. This unique animal model allows for genetically identical individuals housed in controlled environments from their first day of life and, using a custom-designed tracking system, we can monitor their behavior continually throughout development. This design allows us to test whether group-phenotypes—even in the absence of environmental and genetic effects—follow predictable developmental trajectories or instead vary unpredictably during early ontogeny, as might occur due to the complexity of individual-social interactions.

We formed (N = 10) groups of Amazon mollies, with each group consisting of four siblings from the same brood. We recorded these groups continuously during daylight hours (12 hours, at 1 fps) from birth through the first 55 days of life (Figure 2). This period covers early development, prior to sexual maturity (which occurs around 3-4 months of age^37,43^). Because tagging newborn fish for individual identification was not possible, behavioral metrics—although derived from individual positions—were summarized at the group-level. These group-level phenotypes comprised two broad classes: averaged individual-level behaviors (mean group velocity and distance from tank center) and intrinsically social behaviors (inter-individual distance and the within-group behavioral correlations for velocity, distance from tank center, and heading angle). These correlations quantify the average degree to which changes in the behavior of one group member align with the behavior of another, providing a measure of social responsiveness. Behavioral metrics were summarized within one-hour bins (12 bins per group per day). We analyzed developmental changes using Bayesian statistics (MCMCglmm^44^), with hour-of-day and day-since-birth as fixed effects, allowing random intercepts and slopes for each group (meaning the model fits a unique line for each group. See Supplemental Statistics for full model details).

**Figure 2.**
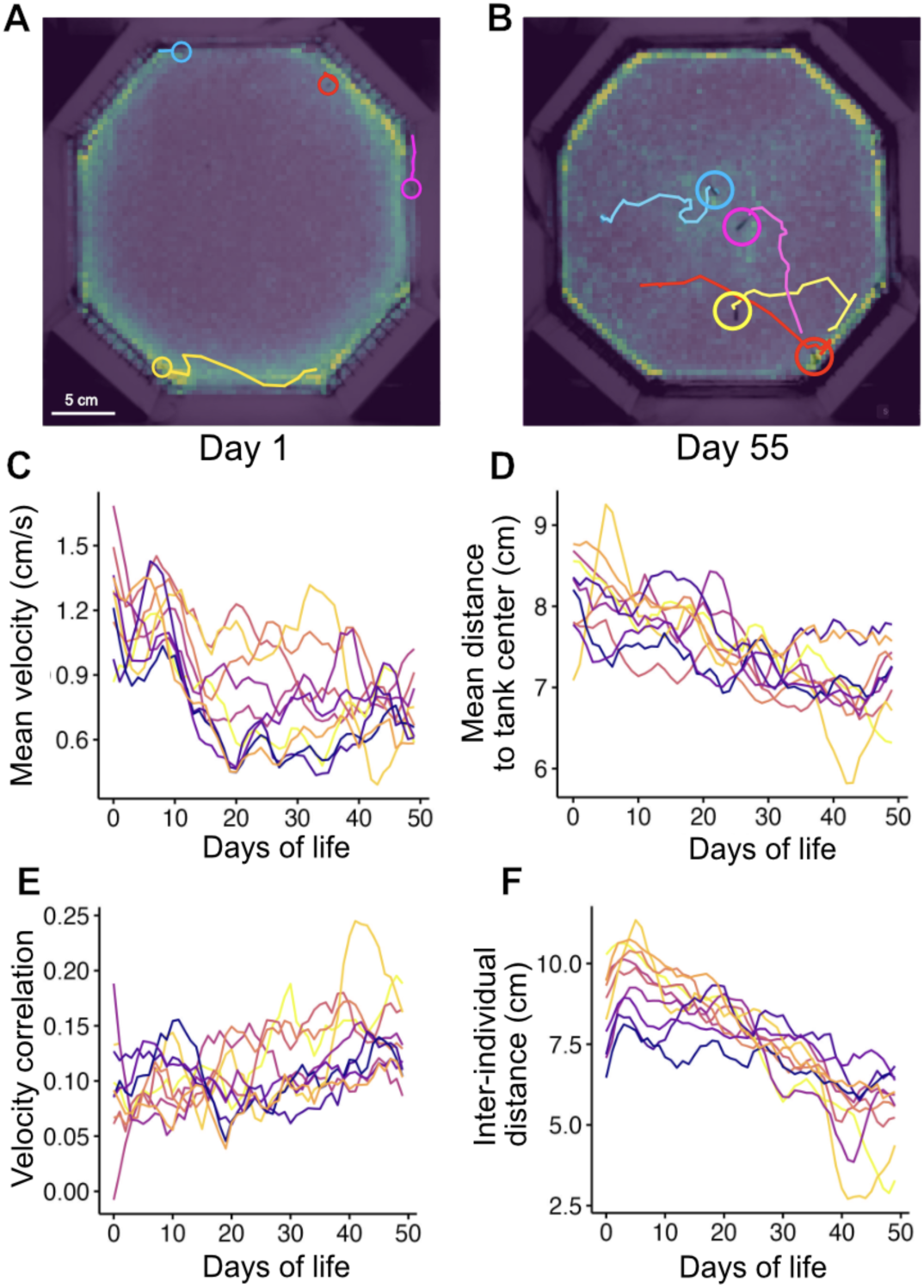
Average group-level behavior changes systematically over time. Panels (A) and (B) show the position of detected fish for a single frame (open circles) and their continuous track from (up to) 1 minute prior, for days 1 and 55 respectively. The overlayed heatmap shows the average space use (brighter colors indicate more time spent in that location). Panels C-F show mean group behavior over time. Each line represents one group, colors correspond to the inter-individual distance at day 1, with highest IID in yellow.

We first characterize average changes in group-level behavioral metrics across the entire 55-day observation period, and then test whether differences among groups exist and how they vary over time. To do so, we use two complementary approaches that quantify the stability and trajectory of group-level behavior across short and longer timescales. First, to test whether and how repeatability changes over time, we use a regression framework to estimate the magnitude of among-group differences for each day of the observation period while accounting for within-group variation across hours within a day. Second, we apply a multivariate approach to estimate among-group correlations across weeks to test how well a group’s behavior at any time point predicts its behavior at later time points.

In regards to average changes in behavior, we observed systematic changes in the mean group-level behavior over the course of development. Across all groups, mean swimming speed decreased over time, similar to prior work in isolated individuals^31^ (effect per day = -0.008, CI: [-0.009 – -0.006], p < .0001; Figure 2C), and fish spent more time in the center of the tank (- 0.025, [-0.030 – -0.188], p < 0.0001, Figure 2D), perhaps reflecting a reduction in stress-like behavior with age. While this could be driven by the arena becoming functionally smaller (relative to body size) as fish grow, group behavior varies over short time windows (Figure 2C-D), whereas body size increases steadily over development (Figure S2), suggesting that growth alone is unlikely to fully explain these patterns.

We also quantified several metrics describing how group members interacted. Groups showed a clear decrease in inter-individual distance (IID; effect: -0.092, [-0.123 – -0.065], p < 0.0001; Figure 2F). In addition, group members became increasingly coordinated, in terms of their correlation in distance from the tank center (0.002, [0.002 – 0.003], p < 0.001), correlation in swimming speed (0.001,[ 0.0003 – 0.0026], p < 0.01, Figure 2E), and the correlation in heading angle (0.001, [0.0007– 0.0018], p < 0.001). Collectively, these changes indicate that individuals became more responsive to one another over development, leading to stronger coupling of movement patterns among group-members.

To determine when interactions among group members emerged, we performed a permutation analysis in which the behavioral time series for each individual track was randomly shifted (by 0 ± 60 minutes), removing temporal alignment among individuals while preserving patterns of individual behavior. This procedure generates the null expectation for group-level metrics if individuals were assumed to behave independently. For all intrinsically social metrics, observed values exceeded those expected by chance (Figure 3C-D, S2B,E). In contrast, for velocity and distance from tank center—behaviors that are not inherently social—observed values did not differ from the shuffled data (Figure 3A-B). Notably, during the first two weeks of life, average inter-individual distance did not differ from chance expectations, suggesting little to no social attraction, but by three weeks of age, cohesion increased, with stronger increases beginning around day 30 (Figure 3D).

**Figure 3.**
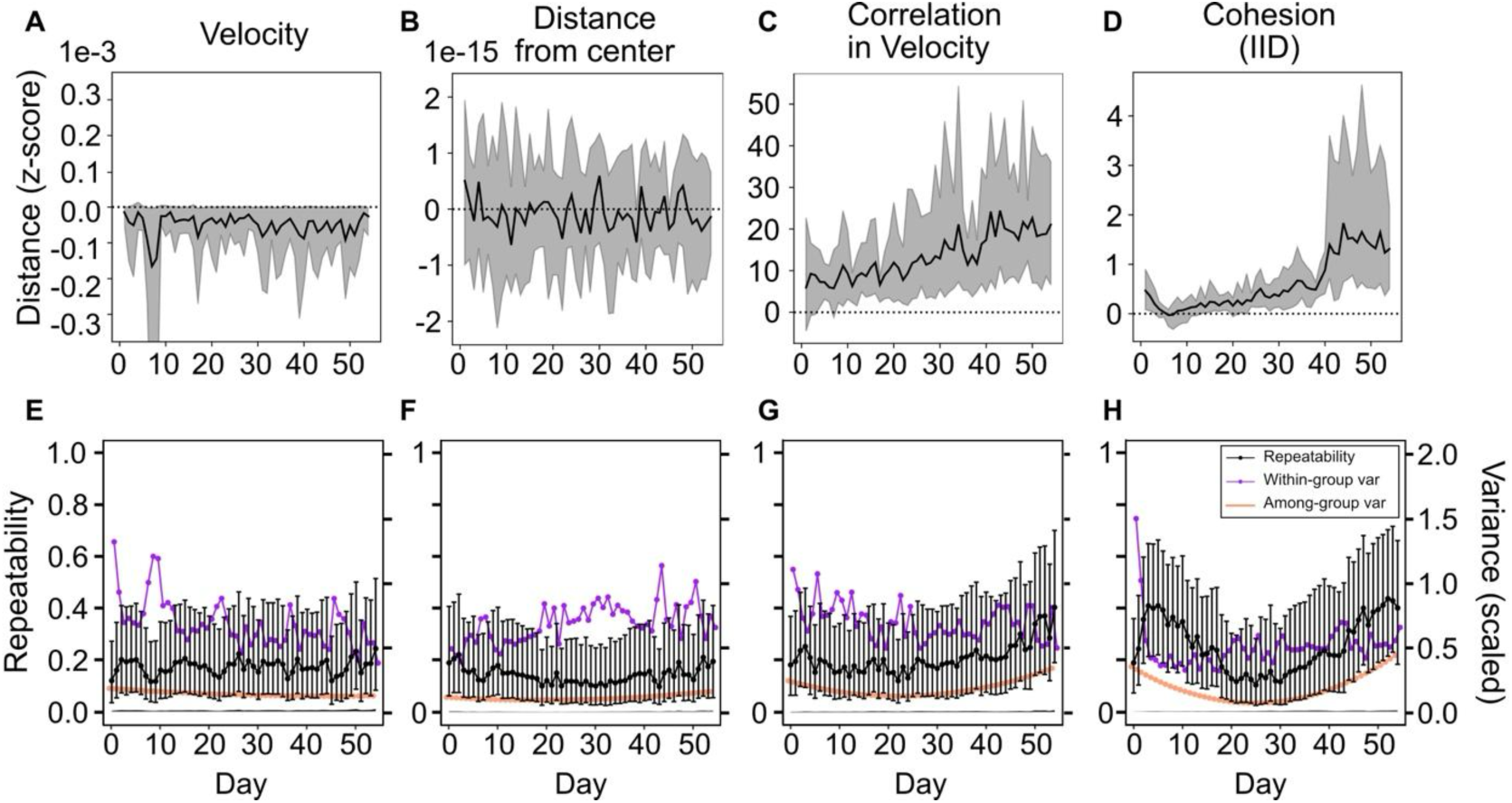
Group-level behavior and repeatability vary over time. The first row shows a permutation analysis for group means of (A) velocity, (B) distance from tank center, (C) pair-wise correlation in velocity, and (D) inter-individual distance. The black line shows the mean difference across time between the behavioral metric and the distribution of individual-shuffled samples, in units of standard deviations for the behavior that day, with the 95% confidence interval shown in gray. The second row shows repeatability (black) and within-group variance (dark-purple) and among-group variance (orange) for (E) velocity, (F) distance from tank center, (G) correlation in velocity, and (H) inter-individual distance. Variance components were scaled to be comparable (right y-axis). The thin black line near 0 shows the shuffled null-distribution of repeatability (STAR-methods).

In addition to describing changes in behavior across groups, we quantified the repeatability (the ratio of among-group variance to within-group variance) of group-level differences over time. Using a random regression approach, we estimated among-group variation in behavior across development. All models yielded significant random slopes among groups over time (Table S1), indicating that the magnitude of among-group differences varied across development. Because within-group variance also appeared to change over time, we modified the conditional repeatability formula from Nakagawa et al^45,46^ to allow day-to-day changes in residual variance^47^. This produces a daily repeatability metric that captures whether groups maintained consistent behavioral differences on short timescales (i.e. within and across days).

This approach revealed repeatable group-level differences from the first day of life (Figure 3E-H), mirroring previous work in isolated newborn molly individuals^31,36,37^. For mean velocity and mean distance from the tank center, repeatability remained relatively stable (Figure 3E-F), consistent both with existing individual data^36^ and with the assumption that individual-level phenotypes are maintained within groups (Figure S1). In contrast, social behaviors showed pronounced temporal structure: repeatability was lower between days 15-35 than in the early and late observation period, resulting in a U-shaped pattern (Figure 3G-H; Supplemental Statistics). This corresponds to groups initially showing clear differences in phenotype, then converging towards similar behavior, before diverging again into distinct group-level phenotypes. This pattern was most evident for inter-individual distance and was supported by a categorical regression contrasting observational phases (early, middle, and late), which revealed phase-dependent differences in repeatability for all behaviors except velocity (Table S2; Figure S3). These differences between social and individualized behavior suggest that the stability of social behavior changes across ontogeny and is sensitive to rapid developmental transitions, broadly aligning with known transitions in sociability during early development^10,37,42^.

The analyses above capture short-term consistency in group behavior. To investigate the long-term stability of behaviors, we used a multivariate Bayesian model on five-day time-binned data, testing whether behavioral differences at a given time (e.g., very early in development) predict the group’s phenotype later in development. We found no correlation for any behavior between group-level phenotype in the first five days of life with the behavioral phenotype exhibited in the last time bin (days 51-55). This contrasts with the known development at the individual level, in which individuals maintain stable rank-order over time^29^, showing that group phenotypes are not simply the average of individual phenotypes (simulated in Figure S1C); nor are they the combination of stable individual phenotypes adjusting towards the group mean, as in some groups of adult animals^2,23,48,49^. Instead, group-level differences in inter-individual distance were only predictable over short time windows, for example, during the first 20 days of life and again during the last 15 days of the experiment, but not in between. Similar patterns exist for the other behaviors measured (Supplemental Statistics). These patterns do not appear to be driven by body size, as group behavior was not generally predicted by mean individual size (Figure S2, Table S3) and behavioral metrics were largely uncorrelated (Supplemental Statistics), suggesting that group-level differences arise from multiple interacting processes rather than a single underlying factor.

## Conclusions

By tracking groups of clonal fish from birth, we show that even with minimal genetic and environmental variation, groups exhibit clear group-level differences, starting from the first day of life, providing a useful null expectation for the early development of collective phenotypes. Previous work has documented short-term differences between groups of adult animals^2,23,48–50^; and group-level divergence occuring over far greater time and distances^18,51,52^. However, no prior work has tracked the emergence of group-level differences over developmental timescales. Although Amazon molly groups were distinct throughout, group-level differences were not stable over time. Rather, groups appear to pass through development transitions, during which their relative behavioral rankings reshuffle. Given that groups lacked significant cohesion for the first 3 weeks, the initial group differences likely reflect intrinsic individual variation, but as individuals mature and social interactions intensify, social feedbacks generate increasingly unstable group-level trajectories^25^, resulting in unpredictable phenotypes.

It is important to note that our observations occurred during early development and in simple, uniform environments. Furthermore, because Amazon mollies display consistent behavioral differences from birth^31,37^, we could not control for phenotypic variation during group formation. Future work can extend this approach to longer timescales and introduce ecological stimuli such as predators or environmental stressors, while reducing baseline group differences further by assembling groups of juvenile individuals with standardized behavioral profiles.

While many questions remain to be explored, this work provides an exceptionally detailed view of how group phenotypes develop and establishes a baseline expectation for how collective phenotypes emerge and reorganize during development, providing a benchmark for future studies dissecting the mechanisms that shape these phenotypes. More broadly, our results show that group-level differences are not reducible to group composition alone: social interactions generate divergent group-level trajectories that are difficult to predict. Identifying the interactions that influence these trajectories will be key to understanding the mechanisms that generate the remarkable diversity of collective behavior across social systems.

## Resource Availability

### Lead contact

Requests for further information and resources should be directed to Ammon Perkes

### Materials availability

This study did not generate new unique reagents

### Data and code availability

All data and code can be found on the paper’s zenodo repository: 10.5281/zenodo.21708571

## Acknowledgments

Funding provided by the Deutsche Forschungsgemeinschaft (BI 1828/2-1 and BI 1828/3-1, to D.B.), under Germany’s Excellence Strategy EXC 2002/1, “Science of Intelligence” project number 390523135 (to M.W. and D.B.), the Alexander von Humboldt Foundation (C.D.), the Zukunftskolleg Institute of Advanced Studies at the University of Konstanz (J.W.J.), the National Science Foundation (IOS-2100625 to K.L.L.; NSF PRFB 2209270 to A.P.), and the National Institute of Health: National Institute of General Medical Sciences of the U.S. National Institutes of Health (1R35GM153309-01 to K.L.L.)

## Declaration of interests

The authors declare no competing interests.

## Author Contributions

Conceptualization, all authors.; Software, A.P., D.B., J.J., and K.L.L.; Formal Analysis, A.P.; Investigation, J.J., C.D., D.B., M.W., and K.L.; Writing – Original Draft, A.P. and K.L.L. ;Writing – Review & Editing, all authors; Supervision, D.B., M.W., and K.L.L.; Funding Acquisition, D.B., M.W., J.J., and K.L.L.

## STAR Methods

### EXPERIMENTAL MODEL AND STUDY PARTICIPANT DETAILS

The Amazon molly (*Poecilia formosa*) is an all-female, naturally clonal, live-bearing fish, that gives birth to genetically identical offspring. It arose from the hybridization of the Atlantic (*P. mexicana*) and sailfin (*P. latipinna*) molly^53–55^. Amazon mollies reproduce through gynogenesis, meaning eggs must be fertilized by the sperm of one of her ancestral (or other Poeciliid) species to stimulate embryogenesis, but the paternal DNA is typically excluded from the egg, resulting in offspring that are genetic clones their mother. Fish begin exhibiting standard social behavior (e.g., fighting) around 4-5 weeks of age^56^, and become sexually mature between 3-5 months of age^35^. Despite their clonal nature, genomic data suggests that their genomes are not showing signs of excessive mutation accumulation, gene conversion or transposable element activity^53^.

All work was performed in compliance with local and federal laws and were approved by the Landesamt fur Gesundheit und Soziales (LaGeSo G-0224/20), the relevant governing body in Berlin, Germany. To obtain experimental animals, we first isolated adult females as potential mothers in individual breeding tanks. The mothers were themselves housed in our lab populations under standardized social and environmental conditions to minimize variation in maternal experience. Each breeding tank contained an artificial plant (a roughly 7 cm ball of green polyester filter fiber) as refuge for newly born offspring. We added two adult Atlantic molly males to each tank for one week to act as potential sperm donors. Amazon mollies typically give birth to broods of around 8-30 offspring. Within broods, individuals are born on the same day as their siblings, usually in the morning. As such, each morning we inspected each breeding tank for offspring and if found, these offspring were carefully netted by trained caretakers in such a way to ensure they were never exposed to the air (which can be stressful for the animals) and immediately placed in our experimental arenas in groups of four siblings (all from the same brood). We initially generated 14 groups of 4. Occasionally, newborn fish escaped through holes in the water outlets in their tanks, died, or our cameras malfunctioned; as such we were able to collect complete data on 10 complete (containing all 4 siblings) groups for the first 55 days of their lives. Altogether, 8 mothers provided the forty offspring that completed the entire 10-week experiment (Supplementary Table S4), meaning two pairs of groups had shared mothers.

### METHOD DETAILS

We used a custom array of tanks (each measuring 27 x 27 cm) to hold the fish throughout the experimental period. Tanks were made from white Perspex, had a water depth of 10cm and were uniformly lit from below using 6500K-LEDs. All tanks were connected to the same filtration system such that water from each tank flowed to the same sump tank, enabling the sharing of chemical cues across all experimental groups and limiting among-tank variation. The tanks were located in an experimental lab with consistent lighting (12h:12h light:dark) and temperature (25 ± 1 °C), maintained by an air conditioning system. All tanks were surrounded by opaque blinds to limit outside disturbances. A standardized amount of powdered flake fish food (TetraMin™) was provided to each tank twice daily, at pseudo-random times to prevent strong time effects. Tanks were cleaned every two weeks by quickly running a gravel-vac along the outside edge, taking care to standardize cleaning across tanks and limit disturbance. Overall, we minimized any potential differences that might occur between tanks by carefully assuring consistent holding tanks, temperature, lighting, feeding, and disturbances.

### QUANTIFICATION AND STATISTICAL ANALYSIS

#### Behavioral tracking

To record the fishes’ behavior, we used Raspberry Pi 3B+ computers attached to a PiNoir camera positioned exactly above the center of each observation tank. We programmed the recordings using the scheduling function of pirecorder^53^ to take a timestamped photo every 1 s during the light period each day, continuously for a period of 10 weeks. Image settings and resolution were optimized to minimize file size while assuring image quality. From the timestamped photos, we then created videos of all recorded images after the experimental observation period. These videos were tracked using Biotracker^54^. Biotracker uses a background subtraction algorithm to provide the x, y coordinates of each fish in each frame. Although the algorithm was able to keep track of individual identities within the small groups quite accurately over short time windows, swaps occurred within the day-long videos particularly due to occlusions (i.e. when one fish swims above another). As such, we focused our analyses on group-level measures that do not rely on individual identities over longer time scales and excluded these putative crosses when relevant (see below).

#### Data processing

We conducted a series of post-processing steps on our tracking to improve track accuracy. To filter out bad detections, we excluded any frames for which the velocity was precisely 0 cm/s (which usually implies the incorrect classification of a stationary object as a fish), or where the velocity was greater than 5.75 cm (200 pixels) per second, which we used as a conservative threshold to exclude any potential switches or bad detection. We also excluded (at the level of pixels) areas of the tank for which there were extremely high rates of detection (a detection density 3 standard deviations greater than the mean), which usually occur due to some stationary feature in the environment being frequently classified as a fish. Finally, because velocity relied on cross-frame consistency, we excluded velocities (but not positions) whenever two fish came within 20 pixels (approximately 7mm) of one another.

Next, we used the final tracking dataset to compute at each frame the displacement, velocity, and heading for each fish, and then computed for every frame the inter-individual distance among the fish, the fishes’ average distance to the tank center, and the individual headings. Heading is calculated based on the velocity vector, and when swimming, is a good approximation of body angle, but at slow speeds, fish often drift sideways or backwards, meaning at these times, the calculated heading may be unrelated to the animal pose. For this reason, we excluded heading (but not velocity) for any frames where the velocity was less than 25 pixels per second (corresponding to 0.71 cm/s).

To quantify the average behavioral measures used for our analyses, we binned each day into 12 1-hour sections, in which we calculated the means for and the mean pair-wise correlations of the speed, distance to tank center, and heading, yielding three individualized behaviors, and three measures of behavioral correlation, as well as spatial cohesion (inter-individual distance). Pair-wise correlation was only performed for frames where both fish had a quantified value (so if a fish was not detected in a given frame—or if we excluded that value due to, for example, low velocity—that frame was necessarily excluded when calculating correlation).

Taking the behaviors and their correlations into account, on each day, for every hour, for every group, we had 6 measures that we analyzed (mean velocity, mean inter-individual velocity correlation, mean inter-individual distance, mean distance from tank center, mean inter-individual correlation of distance from tank center, mean inter-individual correlation of heading angle).

#### Error checking

Our previous work^31^ provided a robust measure of error rates using this tracking system. Briefly, in previous work we manually identified individual fish within photos and then compared these frames with identifications made by Biotracker to determine the rate of errors and whether any obvious bias in error rates occurred across tanks. The overall median error rate over the entire observation period was estimated to be 7%. Because of the addition of group tracking in this dataset, for each group, we manually verified 300 frames from 3 different days (on day 1, day 20, and day 50) for a total of 36,000 frames, checking for bad detections and ambiguous individual labeling within tracklets (which would induce errors in velocity). We found labeling accuracy (in terms of 1 - false positive rate) of 99.9% for individual detections, with a false negative rate of 23%, and 91% for within-tracklet consistency, with 63% of frames assigned to individual tracks. Together, this suggests that any errors introduced by our automated tracking software have minimal if any influence on our behavioral variables and do not affect our interpretation of the results.

#### Statistical analyses

The full details of our analysis can be found in a supplemental R Markdown file, which walks through the analysis step by step, including all results and full model outputs, but we briefly describe the four main statistical approaches used here.

First, we used Bayesian linear mixed models to partition the behavioral variation across different time periods into its among- and within-group components. We ran a separate model for each of the six behavioral metrics (velocity, distance from tank center, correlation in velocity, IID, correlation in distance to tank center, correlation in heading angle) with each model including the fixed effects of day and hour, with group ID as a random slope. Because our question involves change in repeatability over time, we needed to include both a random slope and heterogeneous variance, the latter of which allows the within-group (i.e. residual) variance to vary day by day. We also performed model comparison to assess whether this model structure was truly the best fit for our data. For simplicity, we performed this comparison only on our velocity data. We varied the fixed effects to include Day (of life), Hour, or both, and the random structure with either a) no random effect, or b) random intercepts and/or slopes for groups and c) including or not heterogenous residual variation. In total, we compared 6 models using the Deviance Information Criterion^55^ and found that the Day + Hour, random slope, and heterogenous variance model best described our data (Table S5).

To test whether these group-level behaviors were more synchronized than we would expect by chance, and the extent to which sociality varied over time, we generated null distributions of random behavior by randomly offsetting each individual track by up to one hour (between [-3600,3600] seconds) and repeating this 1000 times. For each iteration, we calculated the distance between the observed value and the shuffled value, dividing it by the standard deviation of the value itself (i.e., for mean group velocity for a given day, the distance between the observed and the shuffled values is divided by the standard deviation of observed velocity for the group for that day). We then calculated the mean of these z-scored distances and the 95% confidence interval by sorting the data and finding the 2.25^th^ and 97.75^th^ percentiles (Figure 3A-D).

Next, to calculate conditional repeatability at each observation day, we extracted the variance components from these models using the formula described^45,46^, modified to allow for heterogenous residual variance^47^. To assess whether repeatability was significantly different from what we would expect by chance, we used a second permutation analysis, this time shuffling the group ID for each hour, to generate a null distribution for comparison (see Figure 3E-H). All models were performed using Markov Chain Monte Carlo estimation with the MCMCglmm package in R v3.6.139. We scaled all data prior to running models and set our models to run 310,000 iterations with a 10,000 burn-in and thinning every 200 iterations, using parameter-expanded priors—which tend to be more robust.

Finally, to investigate how behavioral variance changed over longer time frames, we grouped the dataset into 5-day bins. For each behavior, we then ran a multivariate model with the behavior from each of the bins as the response variables, with the trait as fixed effect and group as a random intercept and estimated the among-group covariance (and correlations) in the chosen behavior across these time bins, along with a 95% credible interval for that correlation. For Figure 4, the points plotted are the estimated intercepts for each group at each time bin, showing the line of best fit.

**Figure 4.**
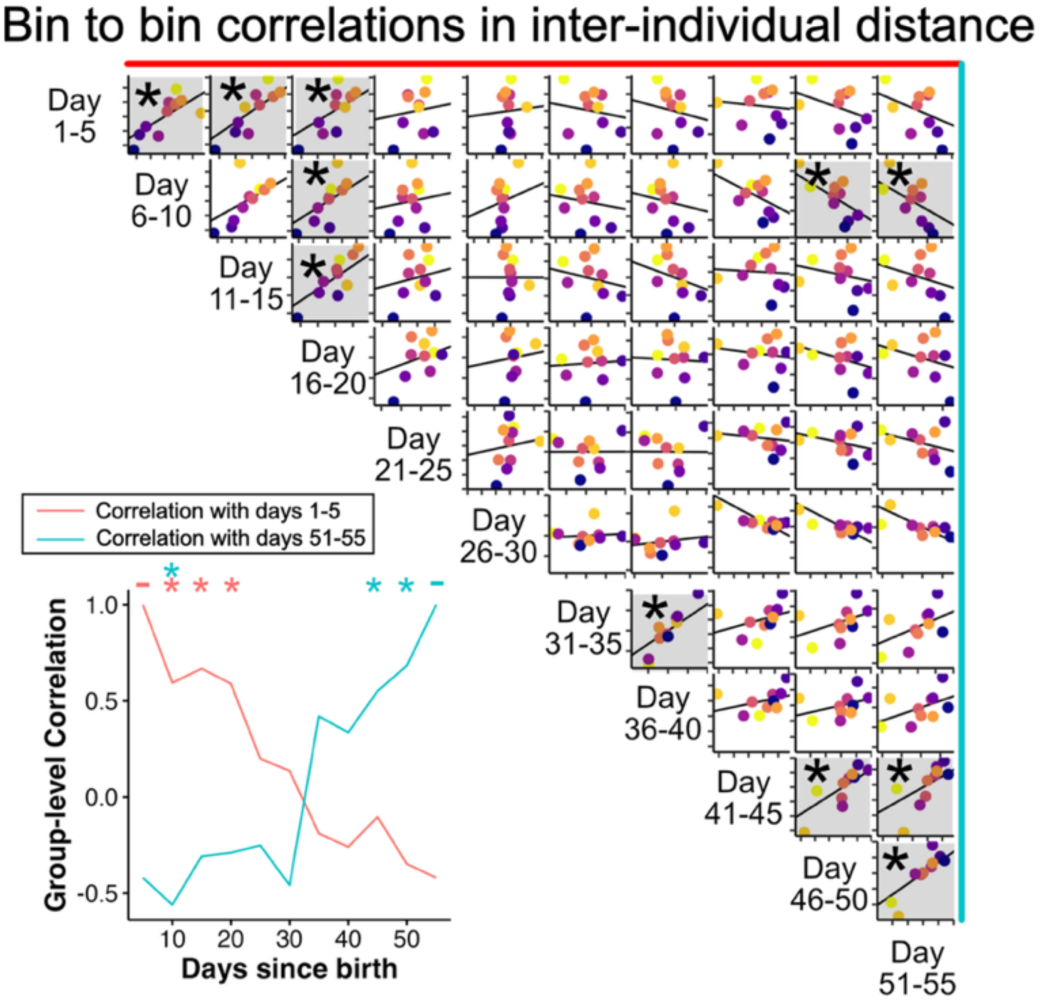
Initial group-level cohesion does not predict eventual group phenotypes. The matrix of plots shows bin-wise correlation for mean inter-individual distance. The top-left-most sub-panel, for example, shows the correlation across groups in inter-individual distance between days 1-5 (y-axis) with days 6-10 (x-axis). Asterisks and shading denote correlations where the 95% credible interval did not overlap with 0. Each dot represents a single group, corresponding to the predicted day-1 IID intercept (highest IID in yellow). The line plots (lower left) show (red line) the estimated correlations for day 1-5 with each subsequent bin and (blue line) correlations between day 51-55 and each preceding bin, corresponding to the topmost and rightmost column, respectively.

In addition to these primary analyses, we performed a few supplemental analyses, which we reference briefly in the main text.

First, to test whether these results could be explained merely from the existence of individual differences over development, we created simulated groups (Figure S1) by averaging velocity across sets of four individuals that were recorded in isolation at the same time as these data (see Laskowski et al, 2022^31^). We then ran the same repeatability and bin-wise correlation as before so that we could establish a baseline expectation of group-level differences.

To assess how repeatability changes over time, we performed a series of categorical regressions between the repeatability of each behavior and the period of the experiment (Table S2), with the periods defined as early (day 1-19), middle (day 20-38), and late (day 39-55).

To assess whether group-level behavior was predicted by the size of the individuals within a group, we first manually annotated the head and tail of fish for 5 frames using SLEAP^57^, in a period of video without crossings. For two groups, we labeled five frames of fish every day for the entire experiment. Then, after confirming that growth was roughly linear, we labeled the remaining groups 5-frames for one day per week. We performed Pearson correlation between IID and mean size at each measured day (day 1, day 5, day 10, etc). We then performed a bivariate regression between size and each behavior, with either the entire data set, the early period (prior to day 30) or late period (after day 30). Because we were interested in ruling out the possibility of correlations, we did not perform any correction for multiple comparisons.

KEY RESOURCES TABLE.

KEY RESOURCES TABLE
| RESOURCE | SOURCE | IDENTIFIER |
| --- | --- | --- |
| **Deposited data** |  |  |
| Processed behavioral data | Zenodo | 10.5281/zenodo.21708571 |
| Sample original videos | Zenodo | 10.5281/zenodo.21708571 |
| Raw tracks | Zenodo | 10.5281/zenodo.21708571 |
| **Software and algorithms** |  |  |
| R 2023.06.0+421 | R foundation | https://www.r-project.org/ |
| Python 3.12.1 | Python software foundation | https://www.python.org/ |
| bioTracker | Mönke et al, 2018<sup>58</sup> | https://www.igb-berlin.de/en/biotracker |
| pirecorder | Jolles et al, 2020<sup>59</sup> | https://github.com/JolleJolles/pirecorder/releases/tag/v3.5.0 |
| **Other** |  |  |
| All code for analysis and pre-processing | Zenodo | 10.5281/zenodo.21708571 |

## Supplemental Figures and Tables

**Figure S1.**
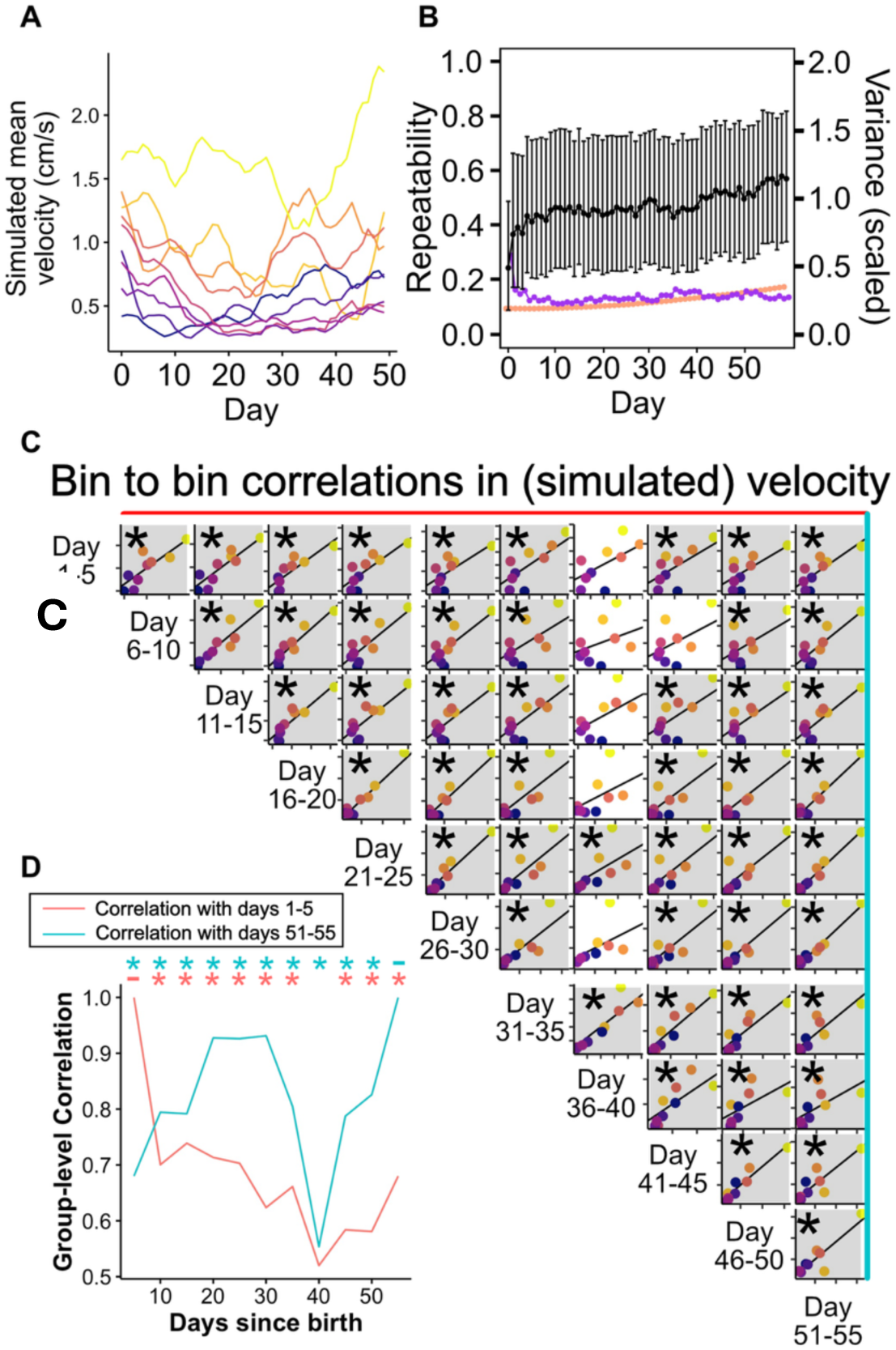
Simulated groups have group-level phenotypes that are distinct but highly consistent over time. (A) and (B) show the mean velocity and repeatability, as in Figure 3, but for groups simulated by taking the mean of velocities from isolated individuals recorded in Laskowski et al, 2022^31^. The large panel of plots (C) shows the bin-wise correlation as in Figure 4, but based on simulated group data, made by taking the average velocity of individuals recorded for Laskowski et al, 2022^31^, with the bottom left panel (D) shows the estimated correlation for every bin with day 1-5 (red) and day 51-55 (blue), which correspond to the top row and far right column, respectively.

**Figure S2.**
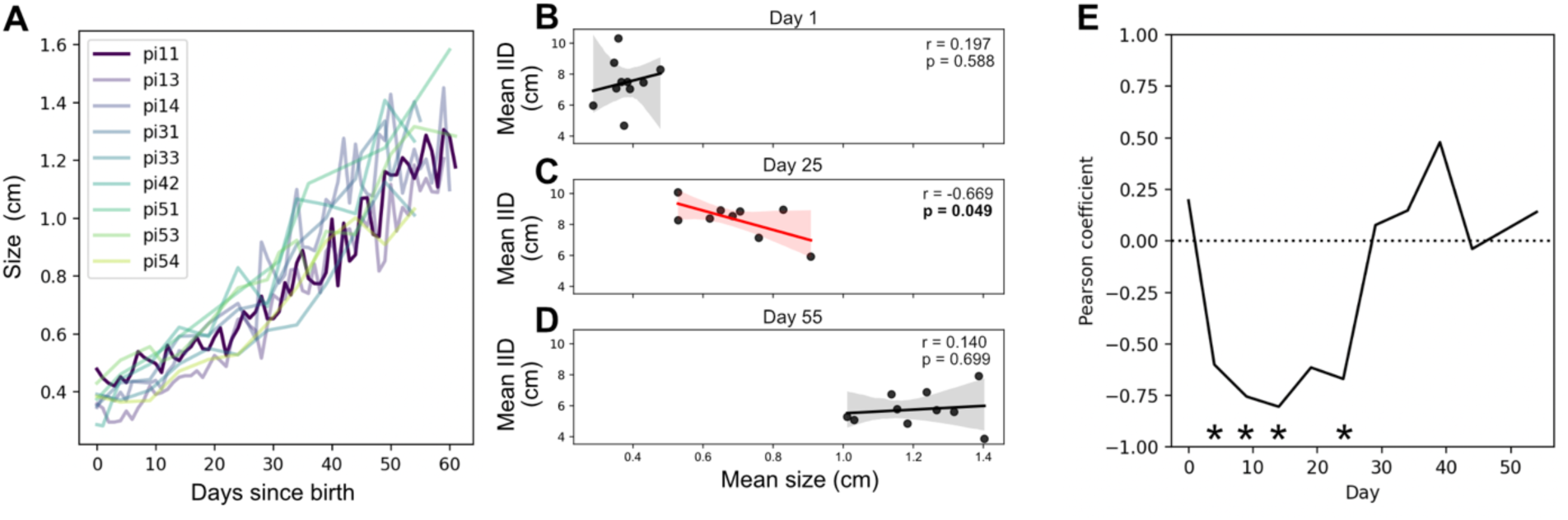
Variation in size predicts group-level cohesion only within a narrow period of development. Panel (A) shows the mean size for each group over time, based on human annotations every fifth day. For two pi’s (dark purple) we annotated the size of individuals for every day. Because annotations were subject to limits in video resolution and human consistency, there is substantial noise in the measurements, whereas body length is expected to be smooth and monotonic. B-D show the best fit line, along with Pearson correlation coefficient, between the mean size of individuals in the group and mean inter-individual distance (our metric of group cohesion) at day 1 (B), day 25 (C) and day 55 (D). Panel (E) shows the Pearson correlation coefficient run at 5-day intervals. Mean size of fish within the group was negatively correlated with group-level cohesion between days 5 and 25, but not afterwards. Asterisks denote time bins for which the credible interval does not overlap 0. These results confirm that the patterns described in Figures 3 and 4 are not merely an artifact of changes in size.

**Figure S3.**
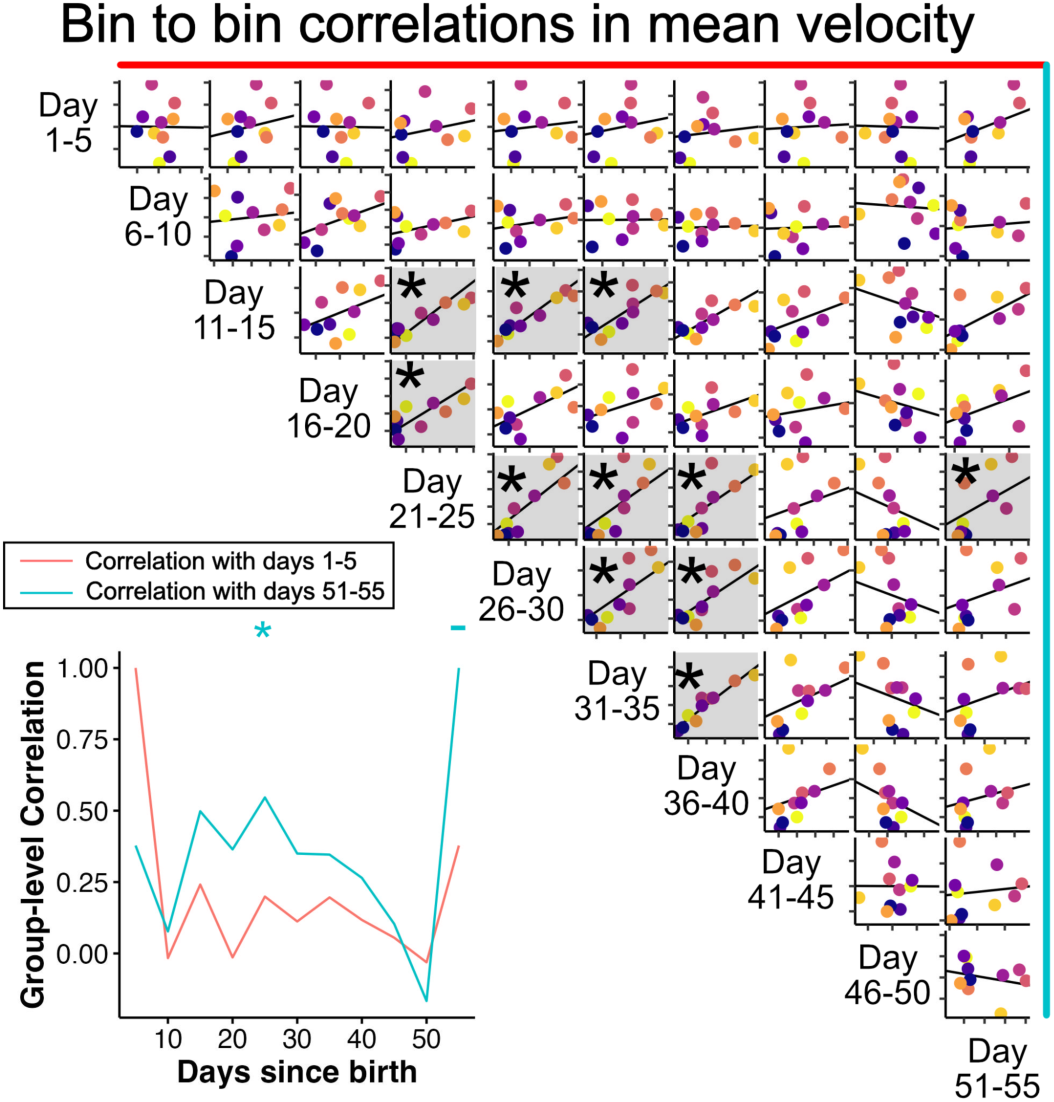
Group-level differences in mean velocity are consistent over time only during mid-development. As in Figure 4, the large matrix of plots shows bin-wise correlation for mean pairwise distance, that is, the top left plot shows the correlation across groups in pairwise distance between days 1-5 (y-axis) with days 6-10 (x-axis). Asterisks and shading denote correlations for which the 95% confidence interval does not overlap with 0. Within the multiple scatter plots, each dot represents a single group, with colors being consistent across plots, and correspond to the predicted intercept of mean pairwise distance (yellow having the greatest distance, i.e., least cohesion, see Figure 2). The bottom left plot shows the estimated correlation for every bin with day 1-5 (red) and day 51-55 (blue), which correspond to the top row and far right column, respectively.

**Table S1.** Model summaries for each behavior. See supplemental statistics for full output.

| Behavior | Day Effect 95% CI<br>(unscaled) | Random Intercept CI | Random Slope CI |
| --- | --- | --- | --- |
| Velocity | (-0.369, -0.246) | (5.435e-02, 4.809e-01) | (6.076e-06, 8.170e-05) |
| Distance to tank<br>center | (-1.012, -0.628) | (0.037, 0.325) | (1.217e-05, 1.671e-04) |
| Correlation of<br>Velocity | (0.001, 0.002) | (0.072, 0.617) | (5.854e-05, 5.851e-04) |
| Inter-individual<br>distance | (-4.191, -2.090) | (0.105, 0.835) | (1.460e-04, 1.088e-03) |
| Correlation of<br>distance to tank<br>center | (3.293e-03, 8.837e-03) | (0.160, 1.333) | (1.757e-04, 1.156e-03) |
| Correlation of<br>heading angle | (3.978e-04, 2.621e-03) | (0.099, 0.696) | (1.690e-04, 1.139e-03) |

**Table S2.** Categorical regression between the repeatability of behavior and the period of experiment (Early: 1-19; Middle: 20-38, Late: 39-55). We report the predicted difference +/-standard error. See Supplemental Statistics PDF for full model output. Bolded values denote statistical significance (all significant values are p<0.0001 or lower)

| Behavior | Middle vs Early | Late vs Early | Adjusted R-squared |
| --- | --- | --- | --- |
| Inter-individual distance | **-0.155 +/- 0.022** | 0.024 +/- 0.22 | **0.586** |
| Velocity | 0.003 +/- 0.009 | 0.008 +/- 0.010 | -0.024 |
| Correlation in velocity | -0.003 +/- 0.014 | **0.086 +/- 0.015** | **0.456** |
| Distance from tank center | **-0.042 +/- 0.008** | -0.003 +/- 0.010 | **0.395** |
| Correlation in distance to tank center | **-0.110 +/- 0.023** | **0.095 +/- 0.024** | **0.567** |
| Correlation in heading angle | **-0.203 +/- 0.019** | **-0.113 +/- 0.020** | **0.681** |

**Table S3.** The correlation between mean size of individuals within a group and each behavioral metric, either based on all of the data, early (prior to Day 30) or Late (Day 30 and onwards). To maximize our ability to detect confounds, correlations were calculated using a bivariate model with only size and the behavior, and there was no correction for multiple comparisons. Bolded values (in this case, only distance in the early category) correspond to p < 0.05.

|  | Inter-individual distance | Velocity | Correlation of velocity | Distance to tank center | Correlation of distance to tank center | Correlation of heading angle |
| --- | --- | --- | --- | --- | --- | --- |
| All | -0.2711966 | 0.07779458 | -0.11264029 | -0.6163554 | -0.3916148 | -0.1286243 |
| Early | **-0.9104420** | -0.0296939 | 0.15072642 | -0.6312208 | 0.2320347 | 0.7446225 |
| Late | 0.3415823 | -0.45618093 | 0.09360457 | 0.1747848 | -0.1023464 | 0.1144187 |

**Table S4.** Overview of fish used. Group number is arbitrary and based on the RaspberryPi camera used to record each group. Note the two pairs of bolded groups, indicating that they came from the same mothers (90.2 and 69.3, respectively).

| Rank<br>(day 1 cohesion) | Group Number | Maternal ID | Start Date | Days recorded |
| --- | --- | --- | --- | --- |
| 1 | 51 | **90.2** | 3/20/18 | 62 |
| 2 | 42 | **69.3** | 3/21/18 | 62 |
| 3 | 53 | **90.2** | 3/20/18 | 62 |
| 4 | 33 | 70.3 | 3/27/18 | 62 |
| 5 | 11 | 80.2 | 4/5/18 | 62 |
| 6 | 14 | 80.1 | 3/1/18 | 62 |
| 7 | 32 | 70.2 | 2/16/18 | 97 |
| 8 | 54 | **69.3** | 3/21/18 | 55 |
| 9 | 31 | 77.2 | 3/30/18 | 62 |
| 10 | 13 | 75.2 | 3/20/18 | 60 |

**Table S5.**
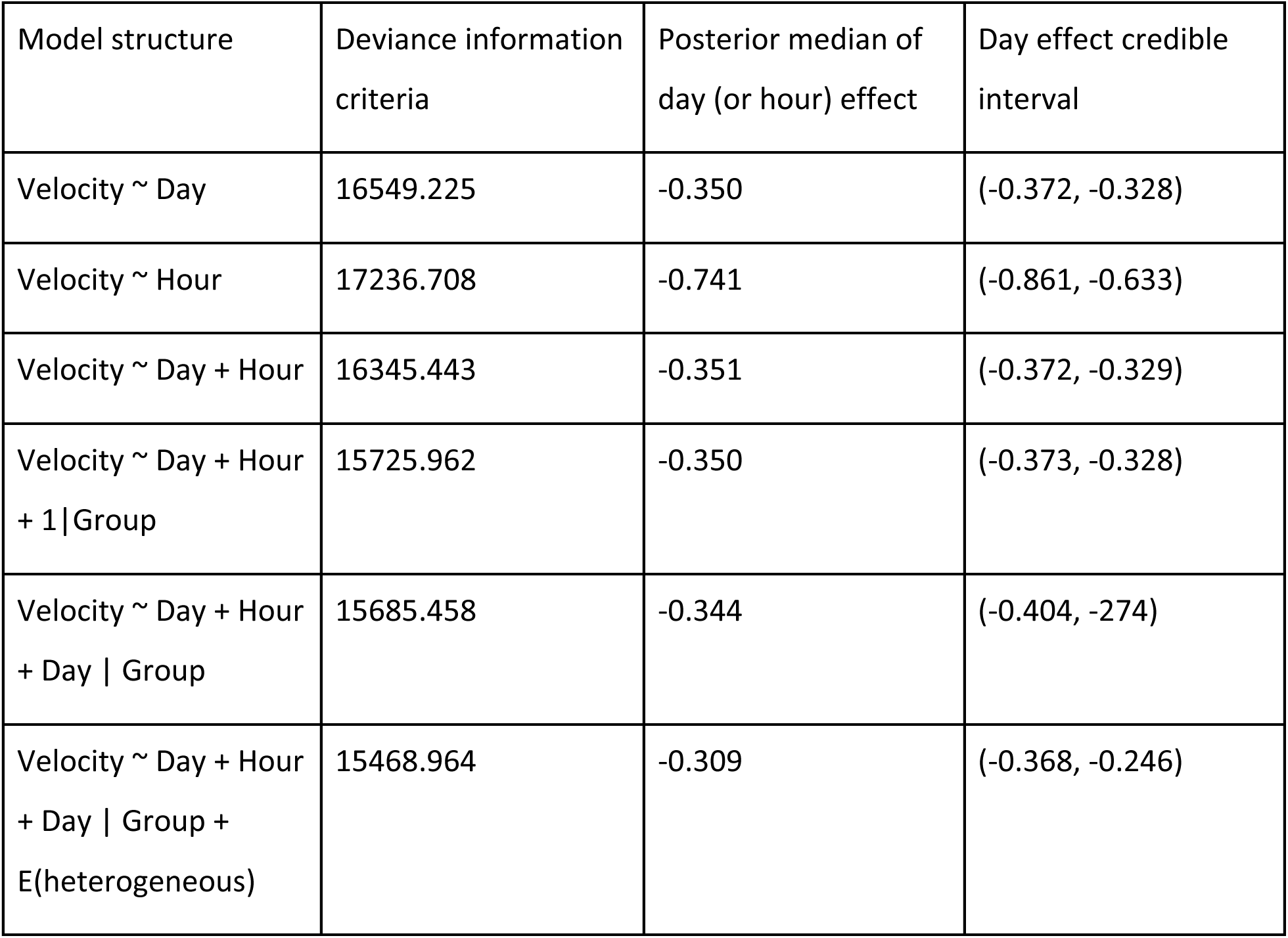
Comparison of various possible models for velocity. The values in the third column show the statistic for the “Day” fixed effect, unless no day is present, in which case it shows statistics for Hour.

## Notes

https://zenodo.org/records/21923853

### Competing Interest Statement

The authors have declared no competing interest.

### Summary of Updates

Based on reviewer comments, the text was updated to improve clarity and made clear both motivations and the limitations of the study. We added a new conceptual Figure 1, showing possible developmental trajectories. We added a new simulation to show the expected group-level differences if we assumed only individual differences. We removed several supplemental figures to focus on the key findings. We updated the Zenodo to contain the entire video dataset.

## Works cited

1. Farine, D.R., Montiglio, P.-O., and Spiegel, O. (2015). From Individuals to Groups and Back: The Evolutionary Implications of Group Phenotypic Composition. Trends Ecol. Evol. 30, 609–621. 10.1016/j.tree.2015.07.005.

2. Jolles, J.W., Boogert, N.J., Sridhar, V.H., Couzin, I.D., and Manica, A. (2017). Consistent Individual Differences Drive Collective Behavior and Group Functioning of Schooling Fish. Curr. Biol. 27, 2862–2868.e7. 10.1016/j.cub.2017.08.004.

3. Viscido, S.V., Parrish, J.K., and Grünbaum, D. (2004). Individual behavior and emergent properties of fish schools: a comparison of observation and theory. Mar. Ecol. Prog. Ser. 273, 239–249. 10.3354/meps273239.

4. Aplin, L.M., Farine, D.R., Mann, R.P., and Sheldon, B.C. (2014). Individual-level personality influences social foraging and collective behaviour in wild birds. Proc. R. Soc. B Biol. Sci. 281, 20141016. 10.1098/rspb.2014.1016.

5. Brown, C., and Irving, E. (2014). Individual personality traits influence group exploration in a feral guppy population. Behav. Ecol. 25, 95–101. 10.1093/beheco/art090.

6. Aplin, L.M., Farine, D.R., Morand-Ferron, J., Cockburn, A., Thornton, A., and Sheldon, B.C. (2015). Experimentally induced innovations lead to persistent culture via conformity in wild birds. Nature 518, 538–541. 10.1038/nature13998.

7. Planas-Sitjà, I., Deneubourg, J.-L., Gibon, C., and Sempo, G. (2015). Group personality during collective decision-making: a multi-level approach. Proc. R. Soc. B Biol. Sci. 282, 20142515. 10.1098/rspb.2014.2515.

8. Smaldino, P.E. (2014). The cultural evolution of emergent group-level traits. Behav. Brain Sci. 37, 243–254. 10.1017/S0140525X13001544.

9. Hinz, R.C., and de Polavieja, G.G. (2017). Ontogeny of collective behavior reveals a simple attraction rule. Proc. Natl. Acad. Sci. 114, 2295–2300. 10.1073/pnas.1616926114.

10. Buske, C., and Gerlai, R. (2011). Shoaling develops with age in Zebrafish (*Danio rerio*). Prog. Neuropsychopharmacol. Biol. Psychiatry 35, 1409–1415. 10.1016/j.pnpbp.2010.09.003.

11. Dreosti, E., Lopes, G., Kampff, A.R., and Wilson, S.W. (2015). Development of social behavior in young zebrafish. Front. Neural Circuits 9, 39. 10.3389/fncir.2015.00039.

12. Harpaz, R., Aspiras, A.C., Chambule, S., Tseng, S., Bind, M.-A., Engert, F., Fishman, M.C., and Bahl, A. (2021). Collective behavior emerges from genetically controlled simple behavioral motifs in zebrafish. Sci. Adv. 7, eabi7460. 10.1126/sciadv.abi7460.

13. Thor, D.H., and Holloway, W.R. (1984). Developmental analyses of social play behavior in juvenile rats. Bull. Psychon. Soc. 22, 587–590. 10.3758/BF03333916.

14. Collias, N.E. (1952). The development of social behavior in birds. The Auk 69, 127–159. 10.2307/4081265.

15. Krause, J., and Ruxton, G.D. (2002). Living in Groups (Oxford University Press).

16. Ford, J.K., Ellis, G.M., Barrett-Lennard, L.G., Morton, A.B., Palm, R.S., and Balcomb III, K.C. (1998). Dietary specialization in two sympatric populations of killer whales (Orcinus orca) in coastal British Columbia and adjacent waters. Can. J. Zool. 76, 1456–1471. 10.1139/z98-089.

17. MacGregor, H.E.A., and Ioannou, C.C. (2021). Collective motion diminishes, but variation between groups emerges, through time in fish shoals. R. Soc. Open Sci. 8, 210655. 10.1098/rsos.210655.

18. van Leeuwen, E.J.C., Cronin, K.A., and Haun, D.B.M. (2018). Population-specific social dynamics in chimpanzees. Proc. Natl. Acad. Sci. 115, 11393–11400. 10.1073/pnas.1722614115.

19. Halfhill, T., Nielsen, T.M., Sundstrom, E., and Weilbaecher, A. (2005). Group Personality Composition and Performance in Military Service Teams. Mil. Psychol. 17, 41–54. 10.1207/s15327876mp1701_4.

20. Milgram, S., Bickman, L., and Berkowitz, L. (1969). Note on the drawing power of crowds of different size. J. Pers. Soc. Psychol. 13, 79–82. 10.1037/h0028070.

21. Clark, C.W., and Mangel, M. (1986). The evolutionary advantages of group foraging. Theor. Popul. Biol. 30, 45–75. 10.1016/0040-5809(86)90024-9.

22. Dornhaus, A., Powell, S., and Bengston, S. (2012). Group Size and Its Effects on Collective Organization. Annu. Rev. Entomol. 57, 123–141. 10.1146/annurev-ento-120710-100604.

23. Dyer, J.R.G., Croft, D.P., Morrell, L.J., and Krause, J. (2009). Shoal composition determines foraging success in the guppy. Behav. Ecol. 20, 165–171. 10.1093/beheco/arn129.

24. Pamminger, T., Scharf, I., Pennings, P.S., and Foitzik, S. (2011). Increased host aggression as an induced defense against slave-making ants. Behav. Ecol. 22, 255– 260. 10.1093/beheco/arq191.

25. Bengston, S.E., and Jandt, J.M. (2014). The development of collective personality: the ontogenetic drivers of behavioral variation across groups. Front. Ecol. Evol. 2. 10.3389/fevo.2014.00081.

26. Gordon, D.M., Dektar, K.N., and Pinter-Wollman, N. (2013). Harvester Ant Colony Variation in Foraging Activity and Response to Humidity. PLOS ONE 8, e63363. 10.1371/journal.pone.0063363.

27. van de Waal, E., Borgeaud, C., and Whiten, A. (2013). Potent Social Learning and Conformity Shape a Wild Primate’s Foraging Decisions. Science 340, 483–485. 10.1126/science.1232769.

28. Muratore, I.B., and Garnier, S. (2023). Ontogeny of collective behaviour. Philos. Trans. R. Soc. B Biol. Sci. 378, 20220065. 10.1098/rstb.2022.0065.

29. Biro, D., Sasaki, T., and Portugal, S.J. (2016). Bringing a Time–Depth Perspective to Collective Animal Behaviour. Trends Ecol. Evol. 31, 550–562. 10.1016/j.tree.2016.03.018.

30. Johnson, J., Sheehy, K., and Laskowski, K.L. (2026). Playing dice with behavior: drivers of stochastic individuality. Trends Ecol. Evol. 41, 55–66. 10.1016/j.tree.2025.10.003.

31. Laskowski, K.L., Bierbach, D., Jolles, J.W., Doran, C., and Wolf, M. (2022). The emergence and development of behavioral individuality in clonal fish. Nat. Commun. 13, 6419. 10.1038/s41467-022-34113-y.

32. Freund, J., Brandmaier, A.M., Lewejohann, L., Kirste, I., Kritzler, M., Krüger, A., Sachser, N., Lindenberger, U., and Kempermann, G. (2013). Emergence of Individuality in Genetically Identical Mice. Science 340, 756–759. 10.1126/science.1235294.

33. Werkhoven, Z., Bravin, A., Skutt-Kakaria, K., Reimers, P., Pallares, L.F., Ayroles, J., and de Bivort, B.L. (2021). The structure of behavioral variation within a genotype. eLife 10, e64988. 10.7554/eLife.64988.

34. Honegger, K., and Bivort, B. de (2018). Stochasticity, individuality and behavior. Curr. Biol. 28, R8–R12. 10.1016/j.cub.2017.11.058.

35. Zipple, M.N., Chang Kuo, D., Meng, X., Reichard, T.M., Guess, K., Vogt, C.C., Moeller, A.H., and Sheehan, M.J. (2025). Competitive social feedback amplifies the role of early life contingency in male mice. Science 387, 81–85. 10.1126/science.adq0579.

36. Bierbach, D., Laskowski, K.L., and Wolf, M. (2017). Behavioural individuality in clonal fish arises despite near-identical rearing conditions. Nat. Commun. 8, 15361. 10.1038/ncomms15361.

37. Scherer, U., Ehlman, S.M., Bierbach, D., Krause, J., and Wolf, M. (2023). Reproductive individuality of clonal fish raised in near-identical environments and its link to early-life behavioral individuality. Nat. Commun. 14, 7652. 10.1038/s41467-023-43069-6.

38. Jolles, J.W., Laskowski, K.L., Boogert, N.J., and Manica, A. (2018). Repeatable group differences in the collective behaviour of stickleback shoals across ecological contexts. Proc. R. Soc. B Biol. Sci. 285, 20172629. 10.1098/rspb.2017.2629.

39. Cantor, M., Maldonado-Chaparro, A.A., Beck, K.B., Brandl, H.B., Carter, G.G., He, P., Hillemann, F., Klarevas-Irby, J.A., Ogino, M., Papageorgiou, D., et al. (2021). The importance of individual-to-society feedbacks in animal ecology and evolution. J. Anim. Ecol. 90, 27–44. 10.1111/1365-2656.13336.

40. Bedbrook, C.N., Nath, R.D., Zhang, L., Linderman, S.W., Brunet, A., and Deisseroth, K. (2026). Lifelong behavioral screen reveals an architecture of vertebrate aging. Science 391, aea9795.

41. Nakayama, S., Masuda, R., and Tanaka, M. (2007). Onsets of schooling behavior and social transmission in chub mackerel Scomber japonicus. Behav. Ecol. Sociobiol. 61, 1383–1390. 10.1007/s00265-007-0368-4.

42. Zada, D., Schulze, L., Yu, J.-H., Tarabishi, P., Napoli, J.L., Milan, J., and Lovett-Barron, M. (2024). Development of neural circuits for social motion perception in schooling fish. Curr. Biol. 34, 3380–3391.e5. 10.1016/j.cub.2024.06.049.

43. Riesch, R., Reznick, D.N., Plath, M., and Schlupp, I. (2016). Sex-specific local life-history adaptation in surface- and cave-dwelling Atlantic mollies (Poecilia mexicana). Sci. Rep. 6, 22968. 10.1038/srep22968.

44. Hadfield, J.D. (2010). MCMC Methods for Multi-Response Generalized Linear Mixed Models: The MCMCglmm R Package. J. Stat. Softw. 33, 1–22. 10.18637/jss.v033.i02.

45. Schielzeth, H., and Nakagawa, S. (2022). Conditional repeatability and the variance explained by reaction norm variation in random slope models. Methods Ecol. Evol. 13, 1214–1223. 10.1111/2041-210X.13856.

46. Nakagawa, S., Westneat, D.F., Mizuno, A., Ajoy, Y.G.A., Dochtermann, N.A., Laskowski, K., Pick, J.L., Réale, D., Williams, C., Wright, J., et al. (2026). Understanding different types of repeatability and intra-class correlation for an analysis of biological variation. 23, 20250545. 10.1098/rsif.2025.0545.

47. Gallagher, J.H., Perkes, A., Chang, C.-C., Chirila, E., Kacevas, K., and Laskowski, K. (2025). Born this way: individuality is seeded before birth and robust to environmental stress. Preprint at EcoEvoRxiv, 10.32942/X2ZH27 https://doi.org/https://doi.org/10.32942/X2ZH27.

48. Knebel, D., Ayali, A., Guershon, M., and Ariel, G. (2019). Intra-versus intergroup variance in collective behavior. Sci. Adv. 5, eaav0695. 10.1126/sciadv.aav0695.

49. Herbert-Read, J.E., Krause, S., Morrell, L.J., Schaerf, T.M., Krause, J., and Ward, A.J.W. (2013). The role of individuality in collective group movement. Proc. R. Soc. B Biol. Sci. 280, 20122564. 10.1098/rspb.2012.2564.

50. Koski, S.E., and Burkart, J.M. (2015). Common marmosets show social plasticity and group-level similarity in personality. Sci. Rep. 5, 8878. 10.1038/srep08878.

51. Weilgart, L., and Whitehead, H. (1997). Group-specific dialects and geographical variation in coda repertoire in South Pacific sperm whales. Behav. Ecol. Sociobiol. 40, 277–285. 10.1007/s002650050343.

52. Aplin, L. (2022). Culture in birds. Curr. Biol. 32, R1136–R1140. 10.1016/j.cub.2022.08.070.

53. Warren, W.C., García-Pérez, R., Xu, S., Lampert, K.P., Chalopin, D., Stöck, M., Loewe, L., Lu, Y., Kuderna, L., Minx, P., et al. (2018). Clonal polymorphism and high heterozygosity in the celibate genome of the Amazon molly. Nat. Ecol. Evol. 2, 669–679. 10.1038/s41559-018-0473-y.

54. Lampert, K. p, and Schartl, M. (2008). The origin and evolution of a unisexual hybrid: Poecilia formosa. Philos. Trans. R. Soc. B Biol. Sci. 363, 2901–2909. 10.1098/rstb.2008.0040.

55. Stöck, M., Lampert, K.P., Möller, D., Schlupp, I., and Schartl, M. (2010). Monophyletic origin of multiple clonal lineages in an asexual fish (Poecilia formosa). Mol. Ecol. 19, 5204–5215. 10.1111/j.1365-294X.2010.04869.x.

56. Laskowski, K.L., Wolf, M., and Bierbach, D. (2016). The making of winners (And losers): How early dominance interactions determine adult social structure in a clonal fish. Proc. R. Soc. B Biol. Sci. 283, 20160183. 10.1098/rspb.2016.0183.

57. Pereira, T.D., Tabris, N., Matsliah, A., Turner, D.M., Li, J., Ravindranath, S., Papadoyannis, E.S., Normand, E., Deutsch, D.S., Wang, Z.Y., et al. (2022). SLEAP: A deep learning system for multi-animal pose tracking. Nat. Methods 19, 486–495. 10.1038/s41592-022-01426-1.

58. Mönck, H.J., Jörg, A., Falkenhausen, T. von, Tanke, J., Wild, B., Dormagen, D., Piotrowski, J., Winklmayr, C., Bierbach, D., and Landgraf, T. (2018). BioTracker: An Open-Source Computer Vision Framework for Visual Animal Tracking. Preprint at arXiv, 10.48550/arXiv.1803.07985 https://doi.org/10.48550/arXiv.1803.07985.

59. Jolles, J.W. (2020). pirecorder: Controlled and automated image and video recording with the raspberry pi. J. Open Source Softw. 5, 2584. 10.21105/joss.02584.

